# *FaDAM3* and *FaDAM4* are candidate genes for the regulation of seasonal dimorphism in cultivated strawberry

**DOI:** 10.1101/2024.12.10.627498

**Authors:** Stephan David, Jielyu Han, Leo F.M. Marcelis, Julian C. Verdonk

## Abstract

Cultivated strawberry (*Fragaria ananassa*) exhibits seasonal dimorphism between the summer and winter seasons. Under the influence of short day conditions, the winter morphology is induced, which includes shorter petioles, smaller and thicker leaves with higher concentrations of chlorophyll and higher density of trichomes. In peach (*Prunus persica*) the Dormancy-Associated MADS-box (*DAM*) gene family is involved in the regulation of winter dormancy, which closely resembles strawberry seasonal dimorphism in terms of timing and environmental regulation. In strawberry, six genes exist with close homology to the peach *DAM* genes. We analyzed the amino acid and coding sequences of these genes, and designated them *FaDAM1-4* and *FaSVP1-2.* We found that there are three pairs of highly similar paralogs within the strawberry genome that exhibit similar expression patterns, although expression levels vary. *FaDAM3* and *FaDAM4* exhibit expression patterns that are consistent with seasonal dimorphism. We conclude that *FaDAM3* and *FaDAM4* are candidate genes for the regulation of seasonal dimorphism in strawberry.

## Introduction

Wild strawberry (*Fragaria vesca*) and cultivated strawberry (*F. ananassa*) exhibit seasonal dimorphism between the summer and winter seasons^1^. Compared to summer morphology, winter morphology is characterized by shorter petioles, smaller, thicker leaves with a higher density of trichomes. While summer leaves senesce and are shed in winter conditions, winter leaves are adapted to the winter cold and drought, and effectively extend the growing season under natural conditions. The winter morphology is induced by exposure to short day conditions (daylengths shorter than 12-14 h) and/or low temperatures^2^. To revert vegetative development from winter morphology to summer morphology, the plant requires exposure to low temperatures (<6°C). The length of the required cold exposure period is known as the chilling requirement. The environmental regulation of strawberry seasonal dimorphism resembles that of winter dormancy in species such as apple (*Malus domestica*) and peach (*Prunus persica*). However, in contrast to these species strawberry does not exhibit cessation of bud growth when dormant, which is why the term semi-dormancy is typically used to describe strawberry dormancy^3,4^.

Winter dormancy consists of two stages: endodormancy and ecodormancy^5^. Endodormancy is maintained by internal signals of the plant^5^. Terminal buds of an apple tree in endodormancy will not grow out, even in favorable environmental conditions. Contrastingly, an endo(semi)dormant strawberry plant will grow new leaves, but exhibit winter morphology^1,6^. After fulfilling the chilling requirement, the plant transitions into ecodormancy. In this stage, growth is inhibited by external conditions (typically cold), but will resume when the plant is moved to favorable conditions^5^. In favorable conditions, an ecodormant apple tree will initiate bud growth, whereas vegetative growth of a strawberry plant will transition to summer morphology.

In seasonal flowering varieties of cultivated strawberry, flower initiation occurs upon extended exposure to short day conditions. These conditions concurrently induce development of the winter morphology^7,8^. Before the plants are cultivated for fruit production, they are therefore stored at low temperatures until the chilling requirement is met. Interestingly, fulfilment of the chilling requirement inhibits subsequent flower initiation^9^. The actual chilling requirement varies between cultivars. Low-chill and high-chill cultivars can be differentiated, with low-chill cultivars typically being destined for climates with relatively short and mild winters, such as in the south of Europe. Importantly, failure to reach the chilling requirement before cultivation can negatively affect vegetative development as well as plant productivity^10^. Chilling for extended amounts of time on the other hand, will reduce the root starch reserves^11^.

It is not known which genes are involved in the regulation of seasonal dimorphism in strawberry. However, since the description of an *evergrowing* mutant in peach, research on the regulation of winter dormancy in fruiting trees from the Rosaceae family has brought to light a family of potential gene candidates^12^. These *evergrowing* trees do not go into endodormancy and thus do not exhibit cessation of terminal bud growth upon exposure to short photoperiods and cold temperatures. This phenotype is caused by a deletion of four genes, named dormancy-associated MADS-box (*DAM*)^12^. These genes are homologs of the MADS-box gene *SHORT VEGETATIVE PHASE* (*SVP*), which delays flowering time in *Arabidopsis thaliana.* MADS-box genes encode transcription factors which contain a DNA binding domain called the MADS-box^13^. The *DAM* genes encode MIKC-type transcription factors, which are a subset of the larger MADS-box family^12^. The abbreviation MIKC refers to the different protein regions that have been identified: the MADS domain (M), a conserved region which is responsible for DNA interaction; the intervening (I) region, which connects the MADS domain and the K domain; the Keratine-like (K) domain, which is involved in protein-protein interactions; and the C-terminal (C) region, which is has been shown to play a role in the stabilization of protein interactions^14^.

Two *SVP* and four *DAM* genes have previously been identified in cultivated strawberry^15^. The similarities between the environmental regulation of winter dormancy in peach and seasonal dimorphism in strawberry, as well as the close evolutionary relationship between these species, make the *SVP* and *DAM* genes likely candidates for regulation of the winter morphology in strawberry. In this work, we explore the phylogenetic relationship between *SVP* and *DAM* genes from strawberry and Rosaceae fruiting trees, as well as the similarities between amino acid sequences of the MIKC motifs. A series of gene expression analyses was performed to investigate which genes are possible gene candidates for the regulation of seasonal dimorphism in strawberry.

## Methods

### Phylogeny and protein alignment

Protein and DNA sequences of strawberry *SVP* and *DAM* genes were obtained from the Genome Database for Rosaceae (GDR)^16^ (Table 1). Sequences of related genes from *Arabidopsis*, peach (*Prunus persica*) and apple (*Malus domestica*) were obtained from Uniprot (uniprot.org) and NCBI (ncbi.nlm.nih.gov). *SVP* and *SVP*-like sequences of *Rosa rugosa* and *Rosa chinensis* were obtained by BLASTN search within the Rosa taxid (taxid:3746) with *FaDAM3* CDS as query sequence (Supplement 1). Amino acid sequences were analyzed for functional domains using Prosite (prosite.expasy.org). Protein alignments and similarity calculation were done in Geneious Prime (version 2024.0.4) with the methods Geneious Alignment, %Similarity (BLOSUM90 with Threshold 0). Phylogenetic tree was constructed in MEGA (version 11.0.13) using the Neighbor-Joining method.

**Table 1.** list of genes that were analyzed in the current study, paired with the entry name in the GDR.

| Gene Name | Entry Name |
| --- | --- |
| *FaSVP1* | Fvb1-2-snap-gene-129.52 |
| *FaSVP2* | Fvb1-4-augustus-gene-107.43 |
| *FaDAM1* | Fvb4-1-augustus-gene-149.40 |
| *FaDAM2* | Fvb4-3-augustus-gene-54.45 |
| *FaDAM3* | Fvb5-2-snap-gene-216.35 |
| *FaDAM4* | Fvb5-3-augustus-gene-39.40 |

### Plant material

All plant material was supplied by Fresh Forward Breeding and Marketing (Huissen, the Netherlands). Two seasonal flowering cultivars were used in this study; ‘Elsanta’, a high-chill cultivar which requires ±1300 chill hours (at ≤7°C) to fully break dormancy, and ‘Fandango’, a low-chill cultivar which requires ±900 chill hours, according to information from the breeding company. The plants were clonally propagated from runners and rooted in plugs of coconut fiber and peat.

### Experiment 1

*F. ananassa* (cv. Fandango) plants were propagated from runners in July 2021 and cultivated outdoors (Huissen, the Netherlands). In October, the plants were moved inside the greenhouse. In the greenhouse supplemental HPS light (SON-T, Philips, the Netherlands) was provided for 10 hours daily and temperatures were kept between 10 and 15°C. On November 8, half of the plants were chilled in darkness at 2-3°C (Chilled), while the other half of the plants remained in the greenhouse (No-chill). On December 18, the Chilled plants were moved out of the chilling conditions back to the greenhouse. On February 16, 2022, eight or six plants were taken from the Chilled and No-chill treatments, respectively, and used for expression analysis. Of each plant, four 1 cm leaf disks were taken from four different leaves and pooled as one sample for gene expression analysis.

### Experiment 2

Young plants of the cultivars Elsanta and Fandango were propagated from runners in the summer of 2022. During the first phase of the experiment, the plants were grown outdoors in plugs positioned in trays under natural photoperiod (ranging from 16h 37m on July 1, to 9h 59m on October 12) and temperature (short day phase). On September 9, September 27, October 12, and October 26, 2022, samples were taken from eight randomly selected plants of each cultivar. The accumulated number of short days (defined as days with a photoperiod of 12 hours or less) on these dates were 0, 2, 17, and 31. Leaves were sampled by taking two 1 cm leaf disks per leaf, of two leaves per plant. The leaf disks of two plants were combined into one biological replicate for a total of four biological replicates per cultivar. For the second phase of the experiment, on November 17, the remaining plants were wrapped in plastic and chilled at 2-3°C in complete darkness (chilling phase). On December 1, December 13, December 23, and January 4 (324, 612, 852, and 1140 chill hours, respectively), plants were taken from the cooling facilities and kept at room temperature for 2 hours, after which leaves samples were taken as described above.

### Experiment 3

Eight Fandango plants from experiment 2 that had been chilled for 1140 hours, were forced (12 hour photoperiod with supplementary far-red light [GreenPower LED Flowering Lamp Gen. 2.1, Philips], average daily temperatures ranging from 13.7 to 15.9°C during the day and 8.7 to 12.5°C at night) in the greenhouse for five weeks, from January 4 until February 8, 2023. At the end of the forcing period, the plants contained winter leaves that had emerged during semi-dormancy (Forced, winter leaf), as well as summer leaves that had emerged during the forcing period after dormancy had been broken by the chilling treatment (Forced, summer leaf). At 13:30, of each plant, three leaf disks were taken from a winter and a summer leaf. Leaf disks of the same leaf type of two plants were combined into one biological replicate for expression analysis. The samples from fully dormant plants from October 26 (Before chilling) and fully chilled plants from January 4 (After chilling) from experiment 2 were included in the expression analysis to provide context to the expression levels. The chlorophyll content of the winter and summer leaves was measured (Force-A, Dualex).

### Confirmation of gene expression

Gene fragments of the *SVP/DAM* genes were amplified with Phusion High Fidelity DNA polymerase (New England Biolabs, Ipswich, MA, United States) using primers designed based on the mRNA sequences obtained from rosaceae.org (Table 1, Table 2). PCR products were analyzed with gel electrophoresis and ligated into the pJET1.2/blunt vector (K1231, Thermo Scientific, Waltham, MA, USA). Resulting vectors were propagated in *E. coli* and the extracted plasmids were sequenced using the pJET1.2 sequencing primers^18,19^. Sequences were uploaded to NCBI.

**Table 2.**
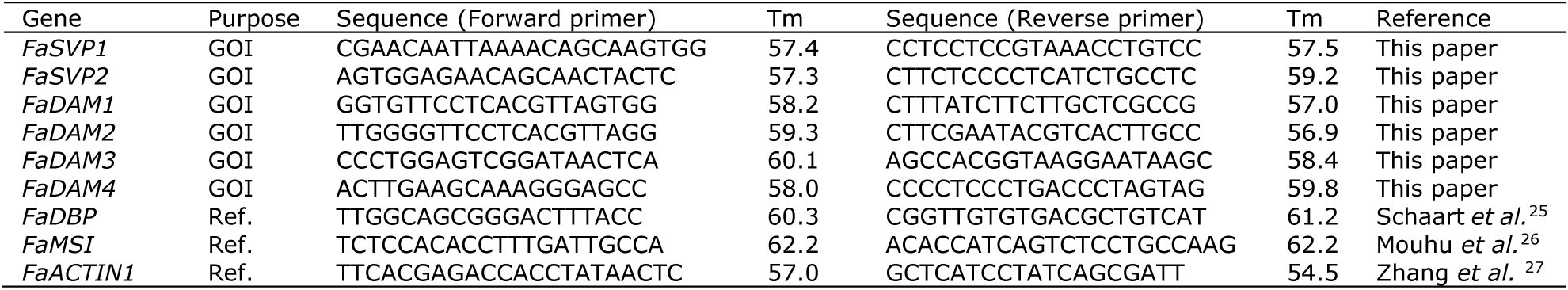
list of primers that were used in expression analysis with qPCR. GOI: gene of interest; Ref.: qPCR reference gene.

### RNA extraction

Leaf material was finely ground in a ball mill grinder (Retsch, MM400, Haan, Germany). RNA was extracted from the plant samples using the CTAB method (adapted from Schultz *et al.* 1994^20^) with lithium chloride precipitation^21^. In a 2 mL microcentrifuge tube, 750 µL extraction buffer (1.4M NaCl, 20 mM EDTA pH8, 100 mM TRIS pH8, 2% CTAB, 2% PVP-40, 1% β-mercaptoethanol) was added to ±50 mg of frozen leaf powder. Samples were incubated for 15 minutes at 65°C. After adding 750 µL chloroform, the samples were mixed and centrifuged at max speed for 5 minutes at RT. The supernatant was transferred to a new microcentrifuge tube containing 500 µL isopropanol, homogenized by inversion and centrifuged at max speed for 5 minutes at RT. The supernatant was removed and pellet was washed with 70% ethanol. The pellet was dissolved in 205 µL MQ and RNA was precipitated by adding 67 µL 8M LiCl and incubating for 30 minutes at −20°C. The RNA was pelleted by centrifugation for 30 minutes at 4°C. Pellet was washed with 70% ethanol, air dried for 5 minutes and dissolved in 50 µL MQ. RNA quantity and quality were assessed by spectrophotometry and gel-electrophoresis.

### cDNA preparation

cDNA was prepared from 200 ng DNAse-treated RNA. DNAse treatment was done using RNase-free DNase (EN0521, Thermo Scientific). cDNA was prepared using the High-Capacity cDNA Reverse Transcription Kit (4368813, Applied Biosystems, Waltham, MA, USA).

### qPCR

Quantitative PCR was performed according to the MIQE^22^, using the iQ SYBR Green Supermix (Bio-Rad, Hercules, CA, USA), and primers from Table 2. A three-step program (initial denaturation for 3 min at 95°C, 40 cycles of 95°C for 10 sec, 30 sec at annealing temperature, and 30 seconds at 72°C, followed by a melt curve from 95 to 55°C) was run on the CFX Thermo Cycler (Bio-Rad). Non-baseline corrected data was analyzed in LinRegPCR^23^ software (version 2021.2) and, where necessary, variation between plates was reduced with Factor_qPCR^24^ (version 2020.0). Samples were normalized for the geometric mean of the expression of three reference genes.

### Statistics

*Student’s t-test was done in Microsoft Excel (two-tailed distribution, homoscedastic). One-way ANOVA was performed in IBM SPSS Statistics (v. 28.0.1.1)*.

## Results

### Protein sequence analysis

The genes discussed in this work were identified by Hytönen and Kurokura^15^. In accordance with the authors, the genes were renamed in the current work according to homology with similar genes in other species (Figure 2). Two of the genes cluster together with Arabidopsis *SVP* and are here referred to as *FaSVP1* and *FaSVP2*. The other four genes are referred to as *FaDAM1-4* (Table 1).

Each protein sequence contains a MADS-box domain, intervening region, K-box domain and C-terminal region (Figure 1). Two MADS-box and K-box domains were found in the FaDAM3 protein sequence obtained from the GDR database (not shown). As it is unlikely that a protein would contain two of these domains, we suspected this to be an error in the database. Alignment between FaDAM3 and its paralog, FaDAM4, showed high similarity between FaDAM4 and the C-terminal half of the FaDAM3 protein. Based on this alignment, we split the FaDAM3 protein and DNA sequences into two halves. We were able to PCR amplify the part corresponding to the C-terminal half, but not the N-terminal half. From here on the C-terminal half is referred to simply as FaDAM3.

**Figure 1.**
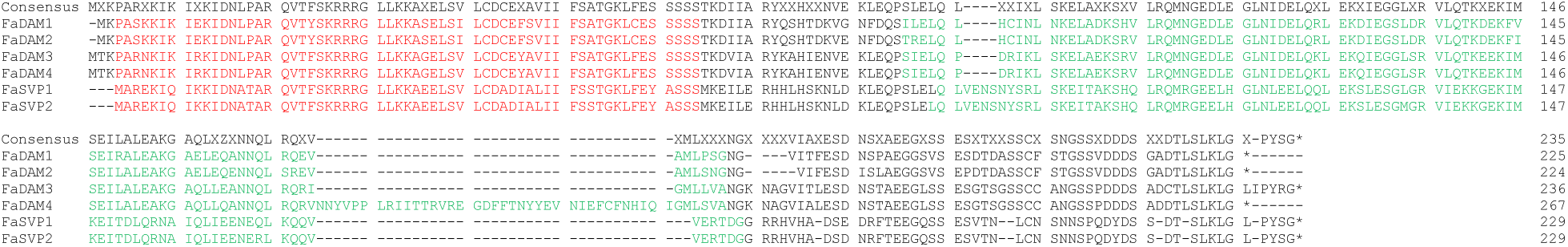
multiple sequence alignment of strawberry SVP and DAM proteins. The consensus is shown on top. The predicted MADS-box and K-box regions are marked in red and green respectively.

### Protein alignment and phylogeny

Protein alignment indicates high sequence identity between the three pairs of paralogs: SVP1 and SPV2 98.2 Percentage Identity (PID); FaDAM1 and FaDAM2 92.4%; FaDAM3 and FaDAM4 84.2%. We investigated similarity between the protein domains (MADS (M) domain, Intervening (I) region, Keratine-like (K) domain, C-terminal (C) region) based on BLOSUM substitution matrix. Alignment of the MADS-box domains showed 100% similarity between each pair of paralogs (Table 3). Similarity of the K-box domains varied from 93% to 100% between the paralogs. Complete similarity was found between all protein regions of FaSVP1 and FaSVP2. The highest dissimilarity between paralogs was found in the C-regions of FaDAM1 and FaDAM2, with 88% similarity. More dissimilarity was found between the protein regions of non-paralogs. In all comparisons the MADS-box domain was the most conserved region of the four.

**Table 3.** Amino acid sequence similarity of the functional motifs and regions of the FaSVP and FaDAM proteins. From left to right, similarity of MADS-box, I-region, K-box, and C-region denoted in % Similarity. Comparisons between the closest paralogs are marked in grey. Similarity was calculated by protein alignment in Geneious Prime (method Geneious alignment, BLOSUM90 with threshold 0).

| % Similarity (MADS-box/I-region/K-box/C-region) |  |  |  |  |  |  |
| --- | --- | --- | --- | --- | --- | --- |
|  | FaSVP1 | FaSVP2 | FaDAM1 | FaDAM2 | FaDAM3 | FaDAM4 |
| FaSVP1 | - | 100 / 100 / 100 / 100 | 85 / 71 / 68 / 55 | 85 / 76 / 68 / 54 | 87 / 76 / 71 / 59 | 87 / 76 / 70 / 60 |
| FaSVP2 | 100 / 100 / 100 / 100 | - | 85 / 71 / 68 / 55 | 85 / 76 / 68 / 54 | 87 / 76 / 71 / 59 | 87 / 76 / 70 / 60 |
| FaDAM1 | 85 / 71 / 68 / 55 | 85 / 71 / 68 / 55 | - | 100 / 95 / 93 / 88 | 93 / 86 / 86 / 78 | 93 / 86 / 81 / 80 |
| FaDAM2 | 85 / 76 / 68 / 54 | 85 / 76 / 68 / 54 | 100 / 95 / 93 / 88 | - | 93 / 90 / 85 / 69 | 93 / 90 / 80 / 71 |
| FaDAM3 | 87 / 76 / 71 / 59 | 87 / 76 / 71 / 59 | 93 / 86 / 86 / 78 | 93 / 90 / 85 / 69 | - | 100 / 100 / 95 / 98 |
| FaDAM4 | 87 / 76 / 70 / 60 | 87 / 76 / 70 / 60 | 93 / 86 / 81 / 80 | 93 / 90 / 80 / 71 | 100 / 100 / 95 / 98 | - |

In phylogenetic analysis, the two strawberry *SVP* genes form a clade together with *AtSVP* and *MdSVPa* and *MdSVPb* (Figure 2). Two genes from the *Rosa* family, one from *R. rugosa* (XM_062150068) and one from *R. chinenesis* (XM_024330395), cluster closely together with *FaSVP1* and *FaSVP2*. We investigated similarity between the functional domains of the SVP proteins based on BLOSUM substitution matrix. The MADS-box motifs of FaSVP1 and FaSVP2 have 100% similarity with MdSVPa and MdSVPb. The K-box motifs of FaSVP1 and FaSVP2 have 97% similarity with MdSVPa and MdSVPb (Supplement 2). These results suggest high functional similarity between the FaSVP and the MdSVP proteins. Strawberry *DAM* genes cluster together in a separate clade with the peach and apple *DAM* genes (Figure 2). Within the *DAM* clade, two separate clades can be distinguished which represent the Rosoideae (strawberry and roses) and the Amygdaloideae (apple and peach).

**Figure 2.**
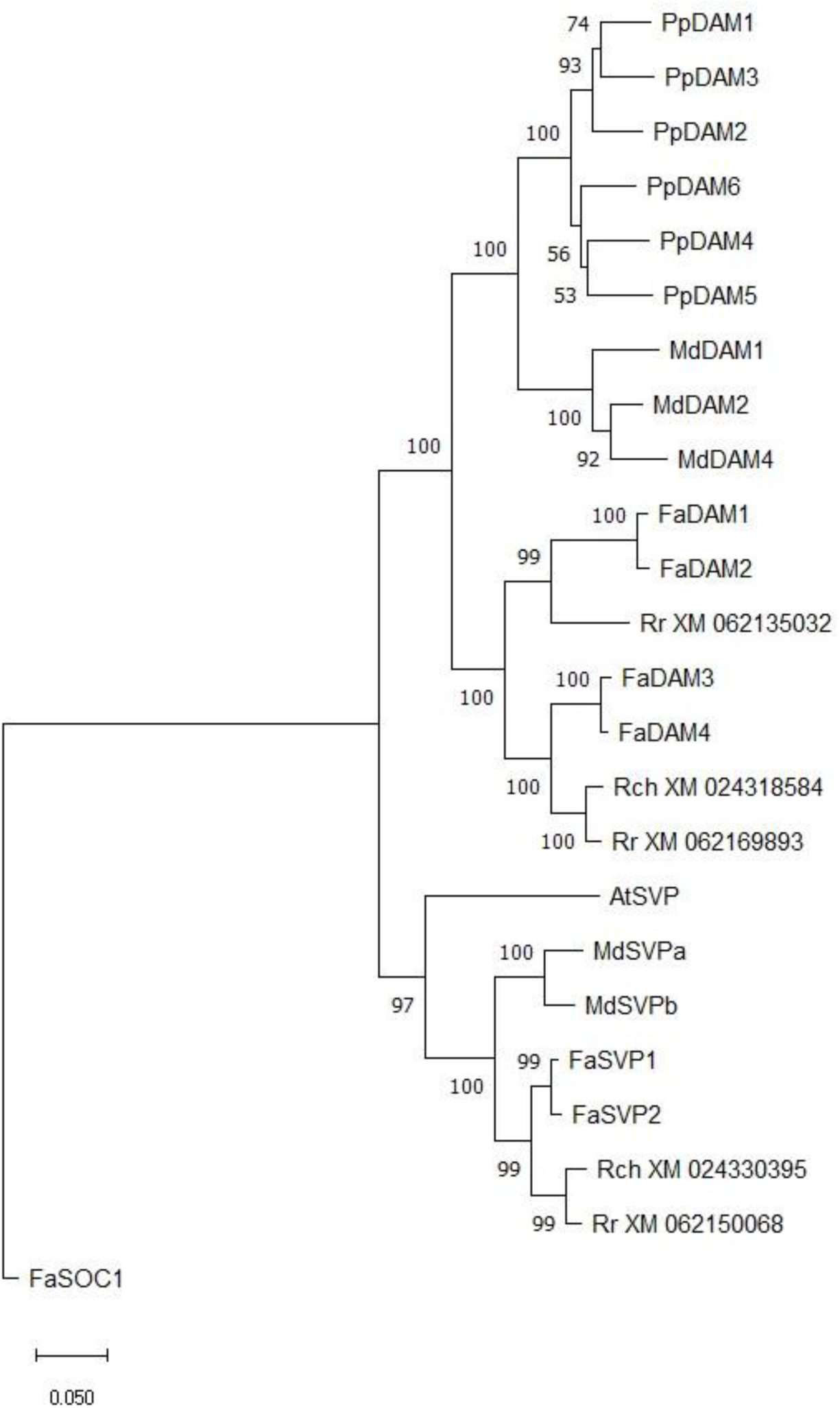
Rooted Neighbor-Joining tree of CDS of SVP and DAM genes. Pp, Prunus persica; Md, Malus domestica; Fa, Fragaria ananassa; Rr, Rosa rugosa; Rch, Rosa chinensis; At, Arabidopsis thaliana. The evolutionary history was inferred using the Neighbor-Joining method^28^. The optimal tree is shown. The percentage of replicate trees in which the associated taxa clustered together in the bootstrap test (1000 replicates) are shown next to the branches^29^. The tree is drawn to scale, with branch lengths in the same units as those of the evolutionary distances used to infer the phylogenetic tree. The evolutionary distances were computed using the p-distance method^30^ and are in the units of the number of base differences per site. This analysis involved 24 nucleotide sequences. Codon positions included were 1st+2nd+3rd+Noncoding. All positions containing gaps and missing data were eliminated (complete deletion option). There were a total of 448 positions in the final dataset. Evolutionary analyses were conducted in MEGA11^31^.

### All members of the SVP and DAM family are expressed in Elsanta

Gene expression analysis of the *SVP* and *DAM* genes is complicated by the high similarity between coding sequences, especially between paralogs (i.e. *FaSVP1 and FaSVP2*, *FaDAM1 and FaDAM2*, and *FaDAM3 and FaDAM4*). For this reason, we wanted to confirm that the individual genes could be amplified with primers that were designed based on the sequences in the GDR.

Gel electrophoresis confirmed that amplicons of the expected sizes had been amplified (data not shown). Sequencing of the cloned PCR products revealed that for all genes, except *FaDAM1*, complete similarity was found between the sequence results and the mRNA entries in Rosaceae.org. In the sequence results of *FaDAM1*, a single mismatch (A176T) was found in the 5’ UTR of the mRNA sequence. This mismatch persisted over cDNA samples from different tissues and likely represents an actual difference in the gene sequence between ‘Elsanta’ and the cultivar used for generation of the data in the Rosaceae database. The mRNA sequences of the *FaSVP* and *FaDAM* genes are included in the supplemental data (Supplement 3).

### DAM3 and DAM4 are upregulated in leaves of dormant plants (Exp. 1)

To investigate which strawberry *SVP* and *DAM* genes are involved in regulation of semi-dormancy, an expression analysis was done on Fandango plants. Half of the plants were grown in short day conditions without chilling (No-chill) and the other half was grown in short day conditions and chilled for four weeks (Chilled). The no-chill plants exhibited the winter phenotype (small, dark leaves, short petioles) whereas the chilled plants exhibited the summer phenotype (large, bright leaves, long petioles) (Figure 3). *FaDAM3* and *FaDAM4* expression in non-chilled plants was 2.3 and 2.4 times higher, respectively, compared to chilled plants (Figure 4). Of the other *SVP*/*DAM* genes, no significant differences in expression were observed between chilled and non-chilled plants.

**Figure 3.**
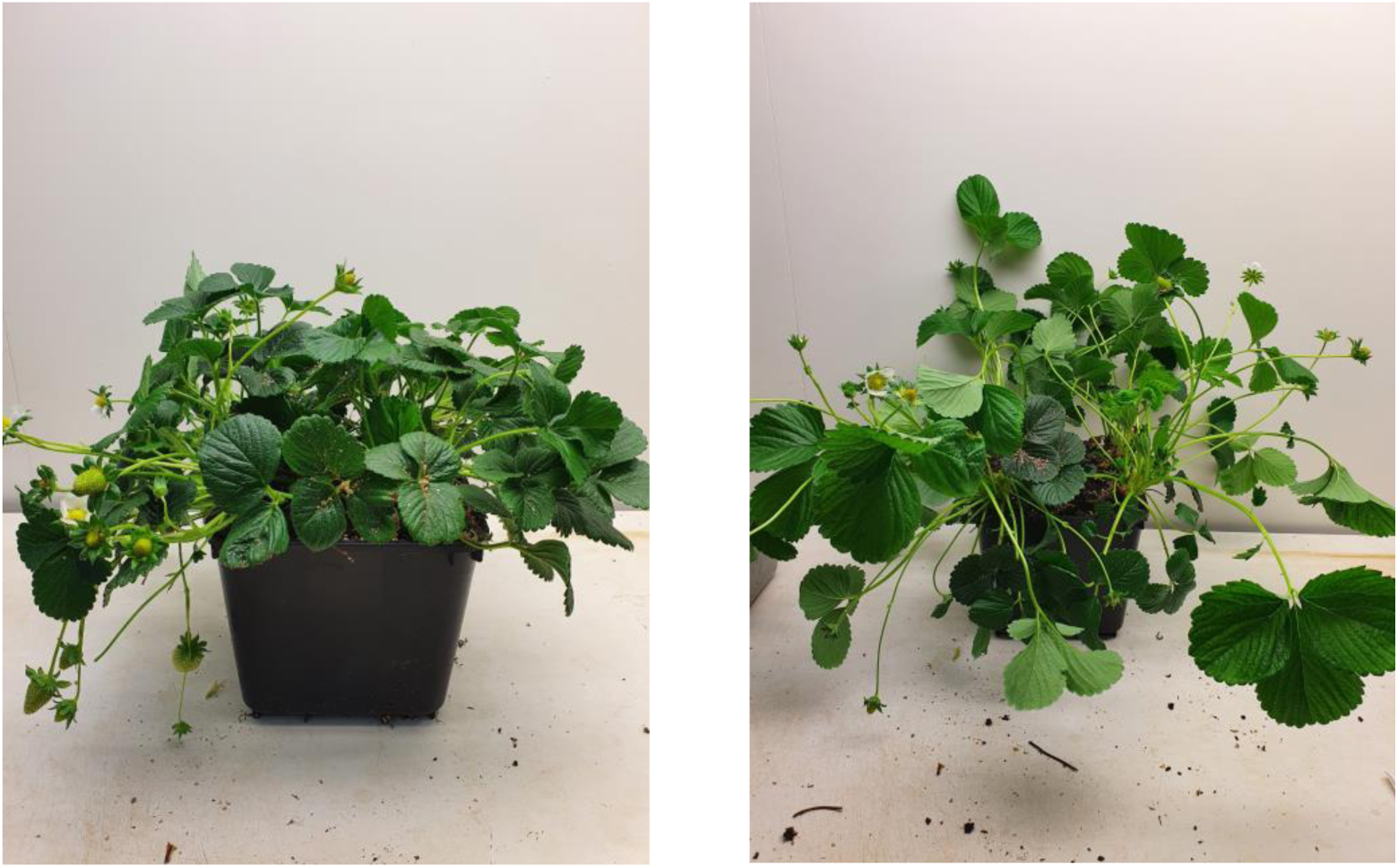
Phenotype of Fandango plants grown in short day conditions without and with chilling. Left: Plants that were grown continuously in short day conditions without chilling (No-chill) and are considered semi-dormant. Right: Plants that were grown in short day conditions and received four weeks of chilling at 2-3°C from November 8 (Chilled) and are considered non-dormant. After chilling they were returned to the greenhouse. Pictures were taken on February 16, 2022. (Exp. 1)

### DAM3 and DAM4 expression increases in short day conditions and decreases during chilling (Exp. 2)

The expression levels of the *DAM* and *SVP* genes were determined at multiple time points during induction (by exposure to short day conditions) and breaking (by chilling) of semi-dormancy (Figure 5, Figure 6). We hypothesized that genes that regulate plant development related to winter-dormancy would be differentially expressed between cultivars with different chilling requirements. Two seasonal flowering cultivars were included in this experiment, Elsanta (a high-chill cultivar) and Fandango (a low-chill cultivar).

**Figure 4.**
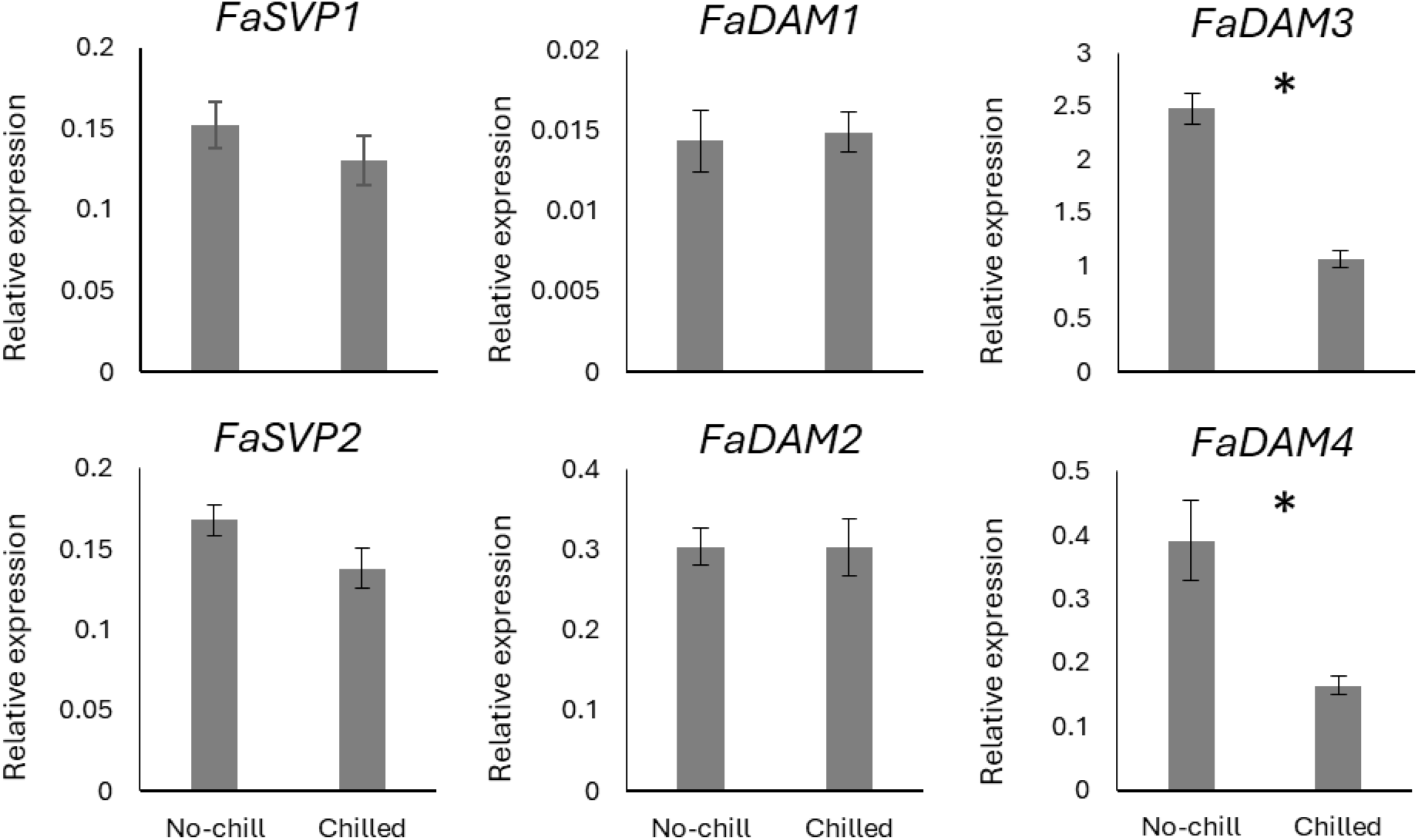
Relative expression of the SVP and DAM genes in strawberry plants of the cultivar Fandango grown in short day conditions without chilling (No-chill), or with 4 weeks chilling at 2-3°C (Chilled). Expression values were determined by qPCR and normalized for the expression of the reference genes FaDBP, FaMSI, and FaACTIN1. Graphs denote the average of eight (FaSVP1 and FaSVP2 Chilled) or six (others) biological replicates ± SEM. Asterisks denote significant differences between no-chill and chilled plants. Statistical significance was determined by t-test (P<0.05). (Exp. 1).

**Figure 5.**
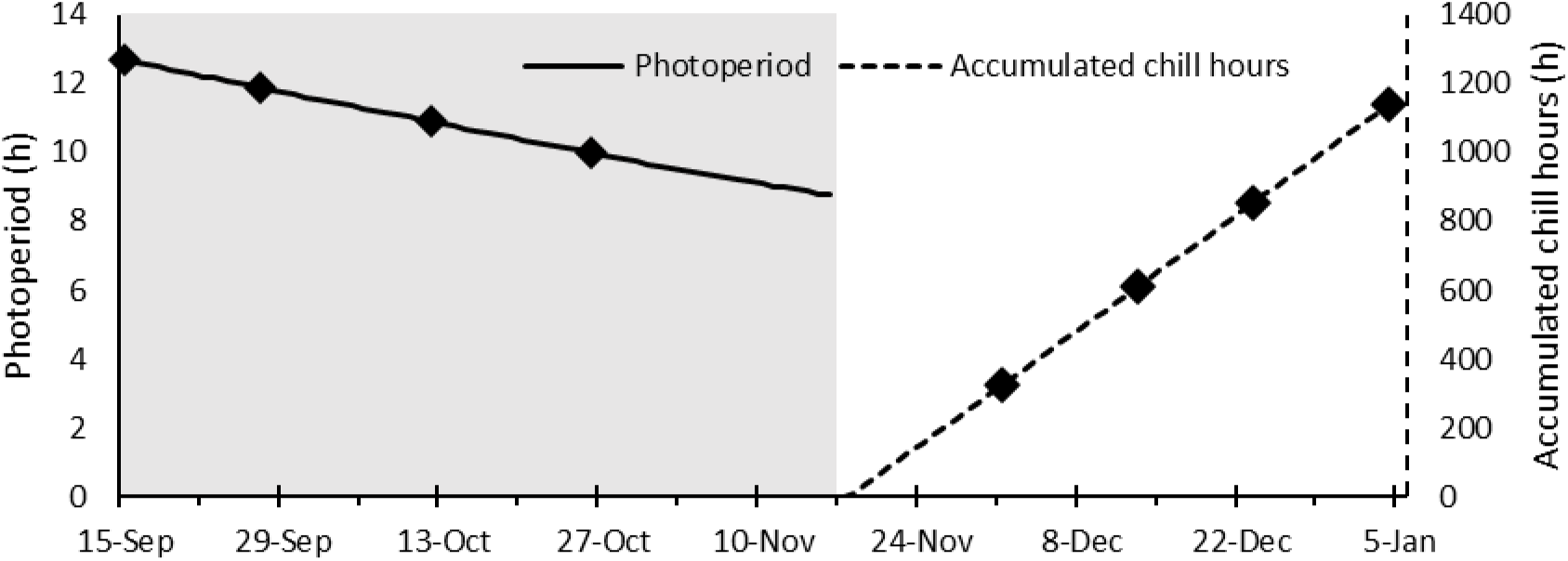
Natural photoperiod (solid line) during the short day phase in the greenhouse (grey area) and accumulated chill hours at 2-3°C in darkness (dashed line) during the chilling phase (white area). Markers denote dates on which leaf tissue samples were taken. (Exp. 2)

**Figure 6.**
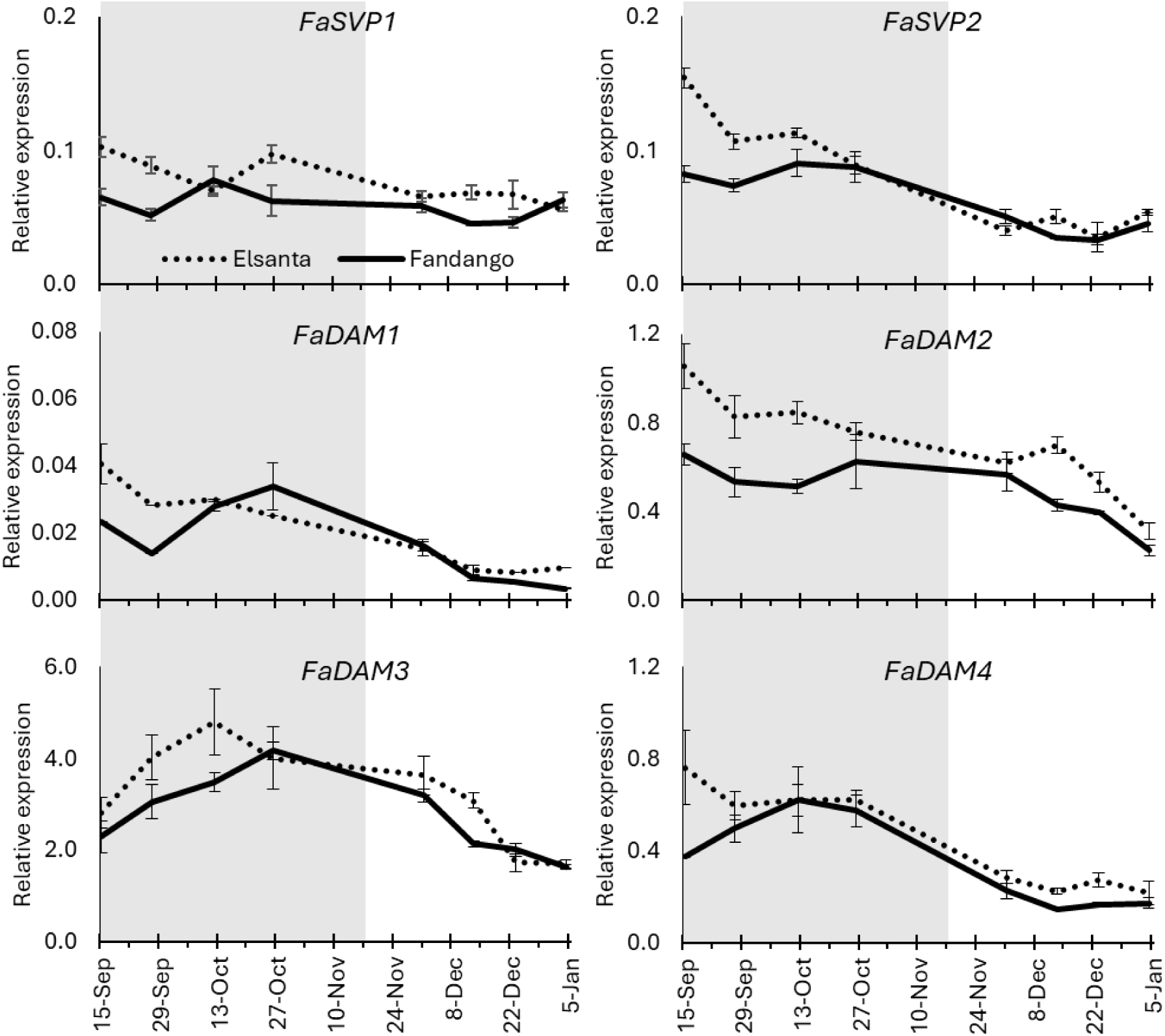
Relative expression of the FaSVP and FaDAM genes in Elsanta and Fandango plants during the short day phase (grey areas) and chilling phase (white areas). Expression values were determined by qPCR and normalized for the expression of the reference genes FaDBP, FaMSI, and FaACTIN1. Graphs denote the average of four biological replicates ± SEM. (Exp. 2).

The relative expression of *FaSVP1* and *FaSVP2* was relatively low in both cultivars throughout the experiment. *FaSVP1* expression was not affected by exposure to short days or chilling. In Fandango, the expression of *FaSVP2* was stable in the short day and the chill phase. However, in the transition from short day conditions to chilling, *FaSVP2* expression decreased. The average expression level over the whole of the chilling phase decreased by 51% compared to the short day phase. In Elsanta, a similar pattern was observed, with the exception that a relatively high expression level was found in the first time-point of the short day phase (Figure 6).

Of the genes that were analyzed in this experiment, *FaDAM1* had the lowest expression levels in both cultivars. The expression was often below the detection rate, resulting in a reduced amount of replicate values and lower reliability of these results. The expression of *FaDAM2* was considerably higher. The *FaDAM2* expression pattern was slightly different between both cultivars. In Elsanta, the expression level decreased over time in both short day as well as in chill conditions, at roughly the same rate. In Fandango, *FaDAM2* expression did decrease in the chilling phase, but was relatively stable during the short day phase (Figure 6).

For both cultivars, *FaDAM3* expression increased during the short day phase and decreased in the chilling phase. Although differences in expression were found on some time points during the short day phase and the chilling phase, the expression pattern of *FaDAM3* was generally found to be the same for both cultivars. The relative expression levels of *FaDAM3* were the highest of all genes that we analyzed in this experiment (Figure 6).

*FaDAM4* expression in Fandango increased during the beginning of the short day phase and stabilized afterwards. After the first 324 hours of chilling, *FaDAM4* expression sharply decreased to below the levels at the start of the experiment, and remained stable afterwards. In Elsanta, a similar *FaDAM4* expression pattern was observed, with the exception that, similar to *FaSVP2*, the expression level was higher in the first time point of the short day phase (Figure 6).

### Expression of FaDAM1 and FaDAM2 is upregulated in summer leaves (Exp. 3)

Because the semi-dormant and non-dormant developmental stages of the plant are characterized by the growth of winter and summer leaves respectively, we sought to investigate whether any of the *FaSVP* or *FaDAM* genes are differentially expressed between these leaf types.

After forcing, the newly emerged summer leaves had lower chlorophyll content compared to the existing winter leaves (Supplement 4). The expression patterns of each pair of paralogs (depicted in the same column, Figure 7) was found to be highly similar. No significant differences in the expression levels of *FaSVP1* or *FaSVP2* were found between the different leaf types. The expression of *FaDAM1* was only successfully measured in one biological replicate of the non-dormant and winter leaf samples. Consequently, no statistical analysis could be done on the expression of this gene. However, of the remaining samples, the *FaDAM1* expression pattern matches that of its paralog, *FaDAM2*. The expression of *FaDAM2* in winter leaves after five weeks forcing was not significantly different from the expression before forcing. *FaDAM2* expression was 2.8 times higher in summer leaves compared to winter leaves (P=0.014). *FaDAM3* expression was significantly reduced in winter leaves after five weeks forcing compared to winter leaves after chilling but before forcing (non-dormant). No differences in *FaDAM3* expression were found between concurrent winter and summer leaves. The expression patterns of *FaDAM4* match those of *FaDAM3*, with the exception that the expression of *FaDAM4* in summer leaves was not found to be significantly different from that in the winter leaves after chilling, but before forcing (Figure 7).

**Figure 7.**
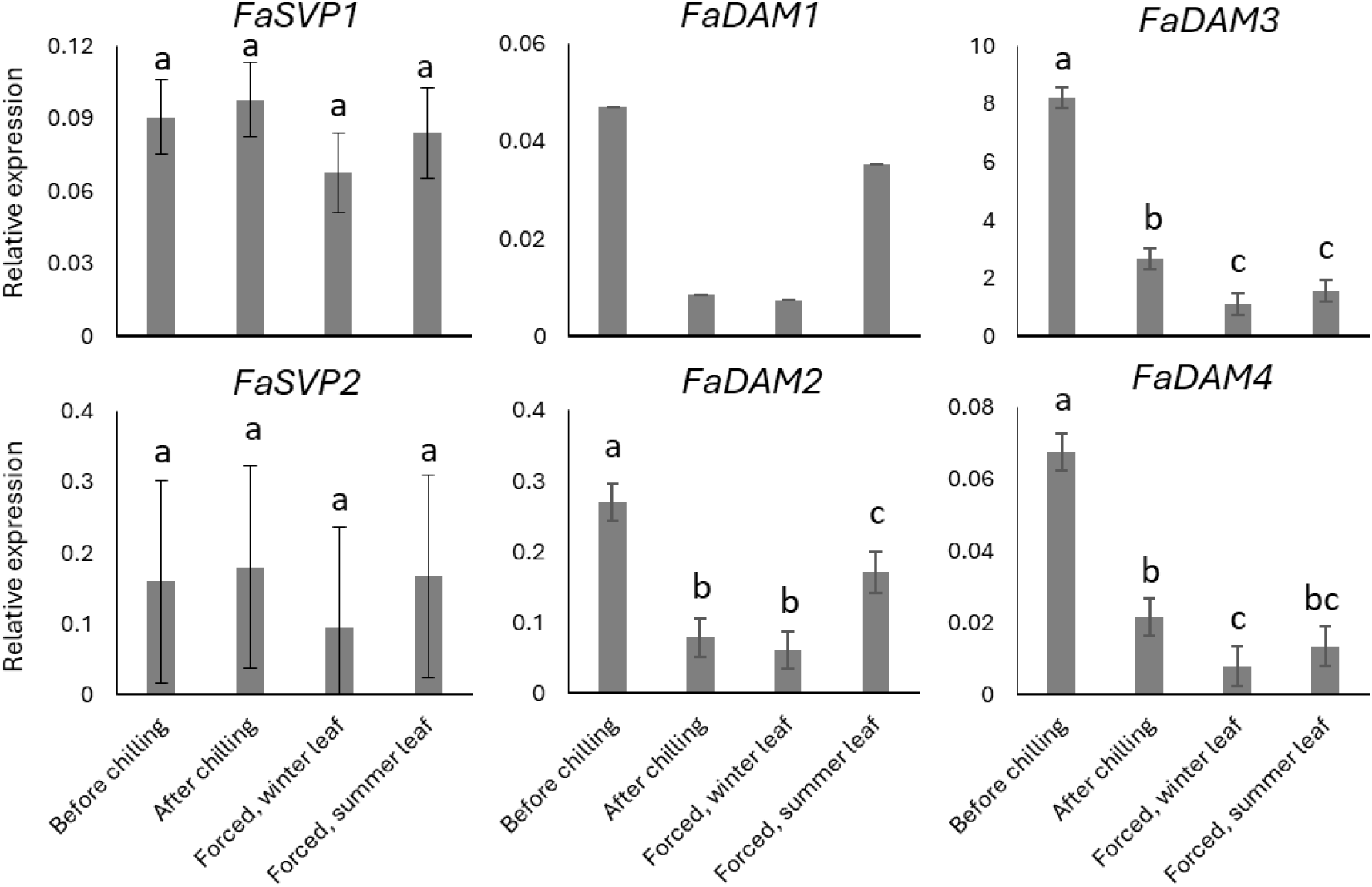
Relative expression of the FaSVP and FaDAM genes in concurrent winter and summer leaves of Fandango plants. Before chilling: plants grown under natural short days until October 26. After chilling: Short day-exposed plants chilled at 2°C in darkness for 1140 hours. Forced, winter leaf: pre-existing winter leaves, emerged during semi-dormancy, of chilled plants grown in the greenhouse for five weeks. Forced, summer leaf: newly emerged summer leaves of chilled plants grown in the greenhouse for five weeks. Expression values were determined by qPCR and normalized for the expression of the reference genes FaDBP, FaMSI, and FaACTIN1. Graphs denote the average ± SEM. Letters indicate significant differences between treatments. Statistical significance was determined by ANOVA. (Exp. 3).

## Discussion

Strawberry plants exhibit seasonal dimorphism and change their vegetative morphology in response to environmental conditions. Vegetative growth continues in the winter, but the leaf morphology is optimized for winter conditions^1^. This differentiates strawberry from peach and other Rosaceae fruiting trees, which undergo complete cessation of bud growth in the winter upon induction of endodormancy. However, similar to the Rosaceae tree species, induction and breaking of semi-dormancy occur through exposure to short days and cold temperatures, respectively^4,32,33^. Although semi-dormancy in strawberry and endodormancy in peach are physiologically very different processes, it is likely that they are regulated similarly, suggesting a role for *DAM* genes in the regulation of seasonal dimorphism in strawberry.

### *DAM* gene duplication in strawberry occurred independently from that in Rosaceae fruiting trees

Phylogenetic analysis shows that the strawberry *DAM* genes cluster together with similar genes from the genus *Rosa*, in a clade separate from those of the Rosaceae fruiting trees (Figure 2). This suggests that, in strawberry, gene duplication of the *DAM* genes occurred after divergence of the Rosaceae subfamilies Rosoideae (which contains *F. ananassa*) and Amygdaloideae (Rosaceae fruiting trees). Because of its small size and comparatively short life cycle, strawberry has the potential to be an interesting model organism to study the *DAM* gene family as a whole. However, because gene duplication and specialization appears to have happened after divergence of the Rosoideae and Amygdaloideae subfamilies, there are limits to what can be extrapolated from one clade to the other.

### Divergence between strawberry SVP and DAM proteins is concentrated in the K-box and C region

Analysis of gene expression and amino acid sequences suggests that all *FaSVP* and *FaDAM* genes that we tested are expressed in strawberry and that translation yields functional proteins. In all protein alignments, the MADS-box region is relatively the most conserved. There is 100% similarity between the M, I, K and C domains of FaSVP1 and FaSVP2 (Table 3). This suggests functional redundancy between these two proteins. Between FaDAM1 and FaDAM2, and FaDAM3 and FaDAM4, most of the dissimilarity was found in the K-box and C regions. The K-box is responsible for protein-protein interaction and the C region has been reported to be involved in the stabilization of K-box interactions^34^ and the formation of protein dimers^35^. The function of MIKC-type MADS-box transcription factors has been reported to change dependent on the formation of protein dimers^36^. Although the high similarity suggests there might be functional overlap between the pairs of SVP and DAM proteins, it is likely that functional divergence has taken place between the paralogs, which may arise from homo- or heterodimerization between members of the SVP and DAM protein families. Furthermore, aside from differences in protein structure, specialization may also occur through differences in temporal or spatial expression.

### *FaDAM3* and *FaDAM4* are likely candidates for the regulation of semi-dormancy in strawberry

*FaDAM3* and *FaDAM4* are upregulated in plants that exhibit signs of semi-dormancy (Figure 3, Figure 4). The short petioles and small leaves of the plants that were not chilled are typical for the winter morphology^1^. The plants used in experiment 1 all had the same age at the time leaf samples were taken. However, because one of the treatments consisted of chilling the plants at 2°C, there is a difference in accumulated growing degree hours between the plants. Because of this, it cannot be said if the chilling treatment has actively reduced *FaDAM3* and *FaDAM4* expression, or simply inhibited its upregulation by slowing down developmental processes. In experiment 2, we demonstrate that *FaDAM3* and *FaDAM4* up and down regulation correlates with exposure to short days and chilling respectively (Figure 6). After the first 324 h of chilling (Dec-1), *FaDAM3* and *FaDAM4* expression had decreased compared to the last datapoint before chilling (Oct-26). This suggests that exposure to cold is actively inhibiting *FaDAM3* and *FaDAM4* expression.

The normalized expression levels of *FaDAM3* were much higher than those of *FaDAM4*. Different levels of *FaDAM3* expression were distinguished during extended exposure to short days and chilling, which is in line with the quantitative nature of the induction and breaking of semi-dormancy^4^. *FaDAM4* expression did decrease upon chilling, but no difference was found between exposure to 324 or 1160 chilling hours. This makes *FaDAM3* a more likely candidate for semi-dormancy regulation, and suggests *FaDAM4* expression might be redundant.

Because we hypothesized that differences in *DAM* expression might be fundamental to the chilling requirement of different cultivars, we included two cultivars with different chilling requirements in this experiment. However, we found no difference between these two cultivars in the rate at which *FaDAM3* or *FaDAM4* expression decreased under chilling conditions. A possible explanation is that differences in chilling requirement manifest at the protein level (e.g. from protein stabilization or interaction), which we did not investigate in the current study. DAM function is known to depend on homo- and hterodimerization of the proteins, and binding-partners are not necessarily other DAM or even SVP proteins^36,37^. It is possible that chilling requirement is fine-tuned by interaction with another protein and the expression thereof. Another possibility is that the difference between high and low chill cultivars arises downstream from *FaDAM3* and *FaDAM4.* In order to further investigate the potential role of *FaDAM3* and *FaDAM4* as master regulators of the winter morphology, we suggest a functional analysis of these two genes by means of gene knock-out and over-expression in a native system.

Flower initiation and semi-dormancy induction occur at photoperiods shorter than about 12-14 h. In the second experiment, we found that the expression of *FaDAM3* and *FaDAM4* was lower after 1160 hours of chilling, than it was at the beginning of the experiment (photoperiod 12h 51m), in both cultivars (Figure 6). If *FaDAM3* and *FaDAM4* expression is indeed induced by exposure to short photoperiods, then it is likely that this induction already took place at photoperiods of more than 12h 51m.

### *FaSVP1* and *FaSVP2* might have no function in strawberry semi-dormancy regulation

Da Silveira Falavigna *et al* (2021) showed diversification between the *MdSVP* and *MdDAM* genes. MdSVP proteins retain the functionality of AtSVP, which inhibits the transition to flowering in Arabidopsis, while MdDAM proteins do not^36^. Phylogeny of *FaDAM* and *FaSVP* suggests that a similar diversification is present in strawberry (Figure 2). Moreover, the high similarity between the MADS-box and K-box sequences of FaSVP1 and FaSVP2 and MdDAM suggests conservation of protein function between these species (Supplement 2). In apple, *MdSVPa* is expressed mostly in the apical bud, and it has been suggested to have a role in bud dormancy^38^. In strawberry, bud growth is not inhibited during semi-dormancy, which suggests that, despite their similarity, these proteins have at least a partially different functionality compared to the apple SVP proteins.

No differences were found in *FaSVP1* expression between dormant and non-dormant plants (Figure 4), or winter and summer leaves (Figure 7). We also found *FaSVP1* expression to be relatively stable during endodormancy up until ecodormancy, with no clear upward or downward trend (Figure 6). This is in accordance with the findings of Da Silveira Falavigna (2021), who also found little variation in the expression of *MdSVPa* and *MdSVPb* during dormancy^36^.

The expression of *FaSVP2* decreased in Elsanta and Fandango after accumulation of 324 chill hours. After this initial decrease, the expression remained stable even when 1140 chill hours were accumulated, breaking endodormancy in both cultivars. The difference in expression pattern between these two genes suggests that, although highly similar in structure, there might not be complete redundancy in function between FaSVP1 and FaSVP2. The expression pattern of *FaSVP2* suggests it may play a role specifically in the induction of endodormancy, but may be of less importance during endodormancy breaking or ecodormancy. However, in all samples included in this study, *FaSVP1* and *FaSVP2* expression has been relatively low (compared to all other genes besides *FaDAM1*). We suggest gene expression analysis of these genes in tissues other than the leaves, specifically the generative and vegetative meristem, as their main function might be outside of leaf tissues. Moreover, the involvement of FaSVP1 and FaSVP2 in processes other than semi-dormancy should be considered.

### Expression of *FaDAM1* and *FaDAM2* varies between leaf types

No clear relationship was found between the expression of *FaDAM1* and *FaDAM2* and exposure to short days or cold temperatures (Figure 6). However, we found differences in *FaDAM1* and *FaDAM2* expression between concurrent winter leaves and leaves that had emerged after chilling and forcing (Figure 7). The lower chlorophyll content of these new suggests that these were summer leaves (Supplement 4)^1^. These genes might play a role in the development and differentiation between these two leaf types. More research is needed however. In each of the experiments *FaDAM1* expression was the lowest of all included genes and could in some cases not be detected by qPCR. Nevertheless, the consistency of the expression pattern between *FaDAM1* and *FaDAM2* suggests that the patterns we found for these genes are accurate.

### Points of consideration

The expression of *DAM* genes in other Rosaceae species is highly seasonal^12,38–40^. Some of these genes have peak expression in summer or autumn and expression levels near zero in the winter months. The leaf samples included in experiment 2 were taken between September 15 and January 4. It is conceivable that some of the genes in this study have peak expression outside of this period, but this has yet to be tested. Furthermore, in apple and peach, not all *DAM* genes are expressed in all tissues^38,39^. Only leaf material was included in the current study. Although we detected mRNA of all genes that we tested, it is possible that some of these genes are (primarily) expressed in other organs, such as the flowers, or vegetative and generative meristem. Finally, *DAM* genes may be involved in developmental processes other than dormancy. *DAM-like* genes are present and expressed in loquat (*Eriobotrya japonica*), a tree species that does not exhibit winter dormancy^40^. Floral initiation is an economically relevant trait in strawberry, which is intimately linked to semi-dormancy. It would be interesting to investigate whether the *SVP*/*DAM* genes play a role in this developmental process. The current study looked into the expression of individual *FaSVP* and *FaDAM* genes. However, it is known that the ability of the proteins to affect transcription of their targets is reliant on homo- and hetero dimerization. To better understand the process of dormancy regulation, and the functionality of FaSVP and FaDAM proteins in general, we suggest an analysis of protein complex formation, and how these complexes affect transcriptional activity.

### Conclusion

We conclude that the expression of *FaDAM3* and *FaDAM4* is consistent with the induction and breaking of semi-dormancy and that these genes are candidates for the regulation of seasonal dimorphism in strawberry. However, further investigation is required to confirm the proposed function of these genes.

## Acknowledgements

This research is part of the TTW Perspectief programme “Sky High”, which is supported by AMS Institute, Bayer, Bosman van Zaal, Certhon, Fresh Forward, Grodan, Growy, Own Greens/Vitroplus, Priva, Signify, Solynta, Unilever, van Bergen Kolpa Architects, and the Dutch Research Council (NWO).

