## Supplements for "*FaDAM3* and *FaDAM4* are candidate genes for the regulation of seasonal dimorphism in cultivated strawberry"

### Supplement 1

Accession numbers of genes that were included in the phylogenetic tree (Figure 2).

| Species name | Name in Figure 2 | NCBI Accession |
|---|---|---|
| Rosa rugosa | Rr_XM_062150068 | XM_062150068 |
| | Rr_XM_062135032 | XM_062135032 |
| | Rr_XM_062169893 | XM_062169893 |
| Rosa chinensis | Rch_XM_024318584 | XM_024318584 |
| | Rch_XM_024330395 | XM_024330395 |
| Malus domestica | MdDAM1 | NM_001328735 |
| | MdDAM2 | NM_001328976 |
| | MdDAM4 | KT582789.1 |
| | MdSVPa | NM_001293986 |
| | MdSVPb | NM_001328882 |
| Prunus persica | PpDAM1 | DQ863253 |
| | PpDAM2 | DQ863255 |
| | PpDAM3 | DQ863257 |
| | PpDAM4 | DQ863250 |
| | PpDAM5 | DQ863251 |
| | PpDAM6 | DQ863252 |
| Fragaria ananassa | FaSOC1 | JQ663534 |

### Supplement 2

Amino acid sequence similarity of the functional motifs and regions of Arabidopsis (At), strawberry (Fa) and apple (Md) SVP proteins. From left to right, numbers denote % Similarity between the MADS-box, I region, K-box, and C region, rounded to the nearest whole number. Similarity was calculated by protein alignment in Geneious Prime (method Geneious alignment, BLOSUM90 with threshold 0).

**% Similarity (MADS-box/I-region/K-box/C-region)**

| | AtSVP | FaSVP1 | FaSVP2 | MdSVPa | MdSVPb |
|---|---|---|---|---|---|
| **AtSVP** | - | 98 / 100 / 90 / 73 | 98 / 100 / 90 / 73 | 98 / 100 / 91 / 67 | 98 / 100 / 87 / 67 |
| **FaSVP1** | 98 / 100 / 90 / 73 | - | 100 / 100 / 100 / 100 | 100 / 100 / 97 / 85 | 100 / 100 / 97 / 83 |
| **FaSVP2** | 98 / 100 / 90 / 73 | 100 / 100 / 100 / 100 | - | 100 / 100 / 97 / 85 | 100 / 100 / 97 / 83 |
| **MdSVPa** | 98 / 100 / 91 / 67 | 100 / 100 / 97 / 85 | 100 / 100 / 97 / 85 | - | 100 / 100 / 96 / 91 |
| **MdSVPb** | 98 / 100 / 87 / 67 | 100 / 100 / 97 / 83 | 100 / 100 / 97 / 83 | 100 / 100 / 96 / 91 | - |

### Supplement 3

*FaSVP* and *FaDAM* mrNA sequences. Lower case: UTR; upper case: CDS; underlined: sequenced. In the case of *FaDAM1* a single base pair in the 5’ UTR has been adjusted based on sequence results (176, A>T). In the case of *FaDAM3*, the sequence which was situated on the 5’ side of the CDS has been left out, because it could not be confirmed by PCR reaction or sequencing.

>FaSVP1_mRNA

tcacaccaccatggttaggatcctattgctgtaactaaaaggcaaaaactttgaaaattccaagtagtttgctttaccataaaaagaccttcccttcctttttcac
gcttccacagacattctgtcattctctctcttcccctccttctcatttcgatccccattggtggcttttgagttttccactccaatccaaccaacacacacagagat
atagagcaccgaaaccaacactgaacctcctcaactgagagggagtttcagttttagggtttgcgcaagattcttagctaaatttggcaaaatggcgccccc
actggacctgtccctctcttattagcctaatttcctgcttttctcacgcataacaacaacttccttgccttttctgaataagaataaagcgggagaagaagatcaa
gaatttgagtgATGGCTCGGGAGAAGATTCAGATCAAGAAGATCGACAACGCCACGGCGAGGCAGGTGACGTTTTCAAAGA
GGAGAAGAGGGCTTTTGAAGAAGGCTGAGGAGCTCTCAGTTCTCTGTGATGCAGATATTGCTCTTATCATCTTCTCTTCAA
CTGGAAAGCTCTTTGAATATGCTAGCTCAAGCATGAAGGAAATCTTAGAAAGGCACCACTTGCACTCCAAAAATCTTGACA
AACTAGAGCAACCATCTCTTGAGTTACAGCTAGTGGAGAACAGCAACTACTCCAGGTTAAGCAAGGAAATAACAGCAAAA
AGCCATCAACTTAGGCAGATGAGGGGAGAAGAACTTCATGGATTAAATTTGGAAGAACTGCAACAGCTGGAGAAGTCCCT
TGAGTCTGGATTGGGCCGTGTGATTGAGAAAAAGGGGGAAAAGATTATGAAAGAGATCACCGACCTTCAAAGAAATGCCA
TTCAGTTGATTGAAGAGAACGAACAATTAAAACAGCAAGTGGTGGAAAGAACTGATGGTGGACGGAGGCATGTTCATGCT
GATTCAGAGGACAGGTTTACGGAGGAGGGTCAGTCATCAGAGTCTGTAACCAATCTATGCAACTCTAATAATTCTCCTCAA
GACTATGACAGCTCAGATACGTCTCTCAAGTTGGGGCTACCATATTCTGGCTGActggaaatggtttgtggtaagttggtgtggttccg
gcctggcctattctgtagggaaattaacttgaagtaaaatgagataaaatactgaactgttaagaaaaaggtgcaagtgatgtattagaacctttgggatag
atctctgtaatgtgtccaagcatcgttaaaagtacttgttttcctagtcactatcacaatgccactcccacaccaatggtgaggcacatctcttctacacaatgcc
acccccacaaatatggtgcacagtttcaacgttgatgtgcttcctcatacttttgattgacggtgcagtctgctttatcctaatcagactcgagaatgatataacg
aagatgaagttcttccatccgatcagatacagttttgtcgttctttgatatctttccagtctagtgtctgaagcacagtccacaatgacttgactccataagtctgt
cttatacttgatggccaattgttggggtagtacatgttgaactattcccattcccggcccggggttgctacttgtgtttggttgaaataaaccagacagaaaatt
ctataccaaataaaatattagaaagttttgaacccgac

>FaSVP2_mRNA

gcaatcgagtgggtacacattgtccttaatagataaaagaaaacctcggtcgtaggcacctaattgtaattaggacacataaatactctccatctgcttggac
ctctacatttgagtcttcacaccaccatggttaggatcctattactgtaactaaaaggcaaaaactttgaaaattccaagtagtttgcttcccttcctttttcacac
ttccacaaacattctgtcattctctctcttcccctccttctcattccgatccccattggtggcttttgagttttccactccaatccaacacacacacagggatatag
agcaccgaaaccaacactgaaccctcctcagctgagaggaagtttcagttttagggtttgcgcaagattcttagctaaatttggcaaaatgtcgcccccactg
gacctgtccctctctcattagcccaatttcctgcttttctcccacataacaacaacttccttgccttttctgagtaagaataaagcgggagaagaagatcaagaa
tctgagtgATGGCTCGGGAGAAGATTCAGATCAAGAAGATCGACAACGCCACGGCGAGGCAGGTGACGTTTTCAAAGAGGA
GAAGAGGGCTTTTGAAGAAGGCTGAGGAGCTCTCAGTTCTCTGTGATGCAGATATTGCTCTTATCATCTTCTCTTCAACTG
GAAAGCTCTTTGAATATGCTAGCTCAAGCATGAAGGAAATCTTAGAAAGGCACCACTTGCACTCCAAAAATCTTGACAAAC
TAGAGCAACCATCTCTTGAGTTACAGCTAGTGGAGAACAGCAACTACTCCAGGTTGAGCAAGGAAATAACAGCAAAAAGC
CATCAACTGAGGCAGATGAGGGGAGAAGAACTTCATGGATTAAATTTGGAAGAACTGCAACAGCTGGAGAAGTCCCTCGA
GTCTGGAATGGGCCGTGTGATCGAGAAAAAGGGGGAAAAGATTATGAAAGAGATCACCGACCTTCAAAGAAATGCCATTC
AGTTGATTGAAGAGAACGAACGATTAAAACAACAAGTGGTGGAAAGAACTGATGGTGGACGGAGGCATGTTCATGCTGAT
TCAGATAACAGGTTTACGGAGGAGGGTCAGTCATCAGAGTCTGTAACCAATCTCTGCAACTCTAATAATTCTCCTCAAGAC
TATGACAGCTCAGATACGTCTCTCAAGTTGGGGCTACCATATTCTGGCTGActggaaatagtttgtggtaaattgtgtggttccggttag
ggagaattaactccaagtaaaatgagatggaatgtaactgaactgttaagaaaaaggtgcaagtgatgtattagaacctttgggttaggtctctgtaatgttt
ccaagcatcgttaatagtacttgttttcctagtcactattacaatgccactcccacattaatggtgaggcaaatcgcttctacacaatgccacccccacaaaaat
ggtgcacagtttcaacgttgatgtacttcctcggtacttttgattgacggtgcagtctactttatctaatcggactcgagaatgataaaacgaagatgaagttct
tccatccgactggtccgagcagatacagatttgtccttctttgacatctttccagtctagtgtctgaagtacagtccacaatgacttgatgaacatctgactccgt
aagtctgtcttatactttatgaccaattgttggggaagtacatgttgaactattcccattcccggcccggggttgcgtacttgtgttaggttgaaataaaccaga
cagaaaattctataccaaataaaatattagaaagttttgaacctgactctttgaacctaccgggccacctctaagaagacgcatggcaattgcttttgatactg
aactttggcatccctcgcattccagtaaatcacctccacaactattctaattgccatctacaacctcccaacctccggccataattatatgattgttggttgatgta
attctatgccacaatcgagcaatgagactagatagatagcaaataggagacgaggcctaccctttgtatctcgttcatg

>FaDAM1_mRNA

gcaaaatacagtagggtgaaggataatgtagaaatatttgacgtctttctttcttaccataaatgggaaacagtcgtttccacagtcgtttccattcttagaag
agcccccaagctccaggcctccagcttggtgttcctcacgttagtggttggtttgatttttgacggaacaataaggatcggagcagatagatacgATGAAG
CCGGCGAGCAAGAAGATAAAGATCGAAAAGATTGACAACTTGCCGGCGAGGCAAGTGACGTATTCGAAGAGGAGAAGA
GGGCTTCTGAAGAAAGCTAGCGAGCTTTCAATTCTCTGTGATTGTGAGTTTTCTGTCATCATCTTTTCTGCTACTGGCAAGC
TCTGTGAGTCCTCCAGCTCCAGTACGAAGGATATCATCGCGAGGTATCAATCGCACACGGACAAGGTGGGAAACTTTGAC
CAGTCAATTCTTGAACTCCAGCTTCATTGCATCAACTTGAATAAAGAACTTGCGGACAAGAGCCACGTGCTAAGGCAGATG
AATGGAGAGGATCTTGAAGGGCTGAACATAGATGAGTTGCAGAGATTGGAGAAAGATATTGAAGGAAGTCTTGACCGTGT
GCTTCAAACTAAGGATGAAAAGTTTGTCAGTGAAATTCGAGCACTTGAGGCAAAGGGAGCTGAATTGGAACAAGCGAATA
ACCAATTGAGGCAGGAGGTAGCGATGTTGCCCAGTGGAAATGGTGTGATCACCTTTGAGTCAGATAACTCGCCTGCTGAA
GGAGGTTCGGTATCAGAGTCCGACACCGATGCCAGCAGCTGCTTTTCTACAGGTTCTTCTGTAGATGATGACTCTGGCGC
TGACACTTTGTCTCTTAAACTTGGGTGAgggagcttgcccggcccttgtcgctttcatcacagagaagtcatgcatgtgtattttgcaacaccgga
atgaaacttgttcatcattttgcactcttctaaatttcaggcttccgtacagtggctaaaactggggagattcagaaggtgaaagtatatatagtatggtttagg
tatttgtaataaaagtacttagagatgccatgcaggttattgtatatatgtatgtattcgatcattttgtgtgtagatgggaagacatggctagtagtgtgagta
acttgccaatgcatttggatggataagatgtagcatccaatatggaaccattagaactaaatgattgtaaacctattcccaaaggttattgtatgaaaaagaa
aaactatatttgcttacgcagccaactaagcatgc

>FaDAM2_mRNA

ctaaactcttttacacgacgatttgttcgggcagcctaaaacgccttgtcgttgtttcccaaaaggaatcaaaatgcaaaatacagctgggtgaaggataatg
tagaaatatttgacgtctttctttctttctttcttaccataaatgggagacagtcgtttccacagtcgtttccattcttagaacagccaccaagctccacgcatcca
gcttggggttcctcacgttaggatcggagcagatagatacgATGAAGCCGGCGAGCAAGAAGATAAAGATCGAAAAGATTGACAACTT
GCCGGCGAGGCAAGTGACGTATTCGAAGAGGAGAAGAGGGCTTCTGAAGAAAGCTAGCGAGCTTTCAATTCTCTGTGAT
TGTGAGTTTTCTGTCATCATCTTTTCTGCTACTGGCAAGCTCTGTGAGTCCTCCAGCTCCAGTACGAAGGATATCATCGCGA
GGTATCAATCGCACACGGACAAGGTGGAAAACTTTGACCAGTCAACTCGTGAACTCCAGCTTCATTGCATCAACTTGAATA
AGGAACTTGCGGACAAGAGCCGCGTGCTAAGGCAGATGAATGGAGAGGATCTTGAAGGGCTGAACATAGATGAGTTGCA
GAGATTGGAGAAAGATATTGAAGGAAGTCTTGACCGTGTGCTTCAAACTAAGGATGAAAAGTTTATCAGTGAAATTCTAGC
ACTTGAGGCAAAGGGAGCTGAATTGGAACAAGAGAATAACCAATTGAGCCGGGAGGTAGCGATGTTGTCCAATGGAAAT
GGTGTCATCTTTGAGTCAGATATCTCGCTTGCTGAAGGAGGTTCGGTATCAGAGCCCGACACCGATGCCAGCAGCTGCTT
TTCGACTGGTTCTTCTGTAGATGATGACTCTGGCGCTGACACCTTATCTCTCAAACTTGGGTGAgggagcttgcccggcccttgtc
gctttcatcacagagaagtcatgcatgtatatttttcaataccggaatgaaacttgttcatcattttgcactcttctaaatttcaggcttccgtaaagtggctaaa
actggggagattcagaaggtgaatgtatatatagtatggtttaggtatttgtaataaaagtacttagagatgccatgcaggttattgtatatatgtatgtattcg
atcattttgtgtgtagatgggaagacatggctagtagtgtgagtatcttgccaatgcatttggatggataagatgtagcatccaatatggaaccattagaact
aaatgattgtaaaccttttctcaaaggttattgtatgaaaaggaaaaactgctaaaaatcaaggtcatgataaaaacttgtagaacaaccatggaaaatggt
agagagaaaacaaactgagctactgggattgataacatcatagttgtacaaaagttggttttcagcttactattttaaaaagggatttttctctatttatactact
tagctattagctattataaacctggaccacaattgggcttatctgttgttcaaatgggccttcatcatcttaacattttcatcatcttaacatccccccctcaaactag
tgattggggcatcaagaactagtttgcagcagagagtattgagccaaaagcttgtaaacagaggaggccaataacttgtctcaaccgggaggaatgagaa
acagtgatcgccatagcttcataggatgagaggcttgtttacaattcagcatatctttttccggagtagaggaatttggaatcggaaaagcttcaattcccttg
acaagagactgctgcaaatgaggataagtaggaactgataatggaagcgcatgtgtaactggcagagccttgacaacatgtgaatgagaagctttcacat
ttgcttgagtagcaatggcaggcggctcctgttgagtgcagatggtagcagacaaagctttaggtcttacaccaacatgaatattcttctctgtatttttggttg
cagtagaaacagttgagcctgccatttcagatacttctgacacattttgcaggatctgaatgtttgatgtcatgtggttcggctttccaagcaccttcgaatctgc
ttgcttggctgctttacacttgttgaattgaggcttgaaatcatctctcctgttgatgcaaccttcattagcaagaaaattaggatagctgatatggccttctgtat
ggctcaggtcactgaacttccttatagaatcttctgcaaaatcaccattctgtgtggaaaacccacagtttgagttattaaaaactggagagtacacttgagag
ctaccattctactgattgaaatttccatttagcattaaagaaatttcaccatctttttgttgtttaaagcttccatgttgctgagaaaatggtgcatttggcaatga
aggaccaacaccattatgcggctgagacatgtatacatccatagtgtcccgaacagcagtgtcttctatagtttctgattcaaacctgtcttttccaccttttcca
attgagaacaactctttcttattctccatttctctcttgctaaaagtagacacatactttctattactcatgaatgcgccagtatgaggaatgcaacttccaccatt
catcattgcatcaatattaaatgaagcttgataaccattagcaatagcaataggcttgacccttttttgaattgttcgcaaagtagattttaactgcataatatga
aactcagaaacaaaagcatattgttgtttaagagtgagccatacctccctcgatgtctggcatccaataacaagggtaagtgcatcagcagaaagtgtggtt
gctatgatacaaatgacaaagctatcttgttgtgtccacttttcataagctccagtgttaacagagagagagtcatgatcagccatctgaacttgggaaggac
aagcatgggaaccatttacaaatttgagtaatccatgaccacgaagatagtgatgcatcaaaaaatgccaagtcgggtaattcgtatcatcaagaagaaca
gagactgcattacaaatttgattcgaggtcgccataacagaactagagcttgaggatgaacaaattttctggaacttgaagaagaatatctacgaacacag
acttgaagaaacgcatgctacgaactcaaaattgaagagaaacaactcccatttcctctctagtaataacagacttcgattttctgatgagtaagactagtaa
ttctacaatggatatttctagggtttttgatttggtacttttgaaaattaaggtttcgaatcgaaattgcagcggagaagtgat

>FaDAM3_mRNA

ATGACGAAGCCCGCGAGGAATAAGATAAAGATCAGGAAGATTGACAACTTGCCTGCGAGGCAGGTGACGTTTTCGAAGA
GGAGGAGAGGGCTTTTGAAGAAAGCTGGTGAGCTTTCGGTGCTCTGTGATTGTGAGTATGCTGTGATCATCTTCTCTGCTA
CTGGGAAGCTCTTCGAGTCCTCCAGCTCCAGCACCAAGGATGTCATTGCAAGGTATAAGGCACACATTGAAAATGTGGAG
AAGTTGGAGCAGCCATCTATTGAGCTCCAGCCTGATCGCATCAAGTTGAGTAAGGAACTTGCTGAGAAGAGCCGTGTGCT
AAGGCAGATGAATGGTGAGGATCTGGAAGGGCTGAATATAGACGAGTTGCAGAAATTGGAGAAGCAGATTGAAGGAGGA
CTTAGCCGTGTACTTCAGACAAAGGAAGAAAAGATTATGAGTGAGATTCTGGCACTTGAAGCAAAGGGAGCCCAGTTGTT
GGAAGCGAACAATCAATTAAGGCAGAGGATAGGGATGCTATTGGTTGCAAATGGAAAGAATGCTGGTGTAATCACCCTGG
AGTCGGATAACTCAACGGCTGAAGAAGGTTTGTCATCGGAGTCTGGCACAAGTGGCAGCAGTTGCTGCGCTAATGGTTCT
TCCCCAGACGATGACTCTGCTGACTGCACTTTATCTCTCAAGCTTGGGCTTATTCCTTACCGTGGCTGAtgagcagggagatca
gaaagtggaagtattttaagcatggcgttgtaataaacgactgtggtcgaagtattttaagaagctatgcagcagggtatttgtatgtatgtctttcttacatg
ggtgcatgtgtgagtgactggtgcttggatgacaaaacataacccgagcccatcgtttgtacaactatatttactgtgtggtgtatgtaaaactgtttttcacg
tttgatttactatgcagcaattttagttacctgcggt

>FaDAM4_mRNA

tttctgaatacctgcccaaaattgggttttactacagtatttgacctcatattcttcccaactctgctcctgcagtacttctggtgacacccgaagcttcagttctgc
accaccaagaaagcaattgagcagctagcagcttacgatgaataagataaggatcgagaagatagagaacctgccggcgaggcaggtgacgttttcgaa
gaagagaaaaggactgttcaagaaagccggagagttatcggttctttgcgacgctgaggttgctattattgtcttctcttctactgacaagctctatgagttctc
caactccagcacggtggatgtcattgcaaggttcaaactgtgcactgaagatgtggaagagggggaccagcagtcacctaatgagctccagacttccatga
tagagagagaacagttaaattattcgaatcaatgattccacttctcgatattagttggtgactgatgagtgcatgaggttgaatatggagcttgcggaaaaga
accacaagctaaggcatatggaggggcaggatctggaagagctgaaaatagatgagttgcagagattggagaatatgattgaaggaggacttagccgc
gtacttcaaactaaggatcagaggattatgagtcagattctggcacttgaaacaaagggagcagagttgacagaagcaaacaacctattaaggcagaggt
taggaacgctatcaaatggagatggaaataaagctagtggcgtcctttcggatccggggatctcaactgatgaagaaggtatgagatcaggatcttgcaca
attgccaccagctgttttagtactggctcttcgagttcgtccatggatgactcctcctctgagtacaccttgtctctcaaacttgggcttcttcctcgctgagctaa
aacatggaaggaaatatggcataattattagtgaataaggtggaattatataatacaccattttggatcgttgtctgtggtgtgtcgaatttgattccaaattcg
aggacatgactagaaggcctgcaggcagatgccagcaaataagatccacggcgtcggtcgtgcaatggaatcatggaagccaggtaccgaactccgga
aagtatcaaatatggaaggggctgccaacttgaagaaaataatactaagatgtgtcctccatcagaacgaaactggcaaggcaaagtgcacgaagttatc
aagccattgatgtagaaagacaaggtcatgtaacgacgttattgagcaactactttttatgcatcataaagagagaggaaagaacatgaaaccctgtata
ttctggttgataaacaagagttcgtgatgagcaaggtgaaacaaccaacgtccttggcccttcgattttcgattttacatcatttcataggtgaatgataacca
acctagcgaagcaaagcaagccaagaacgttgcattggatcgacctaaccctagttcaagtcgtcctgcttgatgtaggtgtgtgtacgctacttaaatatct
atcccttttttttttttttggtcaagtaaatatctatcctgtacaaatcttcatgtcaatatgtacggtatattcatatagtatatattccaaccaaaatagggtggcg
taattagtgacactgaggatgtggggccgaagacatgtataaagcatccccagcacaactagtgtacggacaaatatccaccacatatatatatatggtaa
atgattataccgacacagaattaaattttaaaaaatcaaacccaaaccgacggtggagatcatgtctgttcacgccgaggtatccagatccgtccgatgtgtc
gcaaccgcctccgctcgtctctcccaaagccacaaaaaaaatgcgaaatggcgaccaaaaaaagagagggataagcggggaatatgacgtccttgccctt
ttaccataaatgggaaaaaatatttccattcttgtggtacagccaacaaactcacctccttcagctccaggttcctcagctcccaagaaagctgtgctggctag
atagctatatatagcgATGACAAAGCCCGCGAGGAATAAGATAAAGATCAGGAAGATTGACAACTTGCCGGCGAGGCAGGTGA
CGTTTTCGAAGAGGAGGAGAGGGCTTTTGAAGAAAGCTGGAGAGCTTTCGGTGCTCTGTGATTGTGAGTATGCTGTGATC
ATCTTCTCTGCTACTGGAAAGCTCTTCGAGTCCTCCAGCTCCAGCACCAAGGATGTCATTGCAAGGTATAAGGCACACATT
GAAAATGTGGAGAAGTTGGAGCAGCCATCTATTGAGCTCCAGCCTGATCGCATCAAGTTGAGCAAGGAACTTGCTGAGAA
GAGCCGCGTGCTAAGGCAGATGAATGGTGAGGATCTGGAAGGGCTGAATATAGACGAGTTGCAGAAATTGGAGAAGCAG
ATTGAAGGAGGACTTAGCCGTGTACTTCAGACAAAGGAAGAAAAGATTATGAGTGAGATTCTGGCACTTGAAGCAAAGGG
AGCCCAGTTGTTGCAAGCGAACAATCAATTAAGGCAGAGGGTAAACAACTATGTACCACCGTTGCGTATTATTACTACTAG
GGTCAGGGAGGGGGATTTCTTTACTAATTACTACGAGGTGAATATAGAGTTTTGCTTTAATCACATACAGATAGGGATGCTA
TCGGTTGCAAATGGGAAGAATGCTGGTGTAATCGCCCTGGAGTCGGATAACTCAACGGCCGAAGAAGGTTTGTCATCGGA
GTCTGGCACAAGTGGCAGCAGTTGCTGCGCTAATGGTTCTTCCCCAGACGATGACTCTGCTGACGACACTTTATCTCTCAA
GCTTGGGTGA

### Supplement 4

Chlorophyll content of concurrent winter and summer leaves of Fandango plants. (Exp. 3)

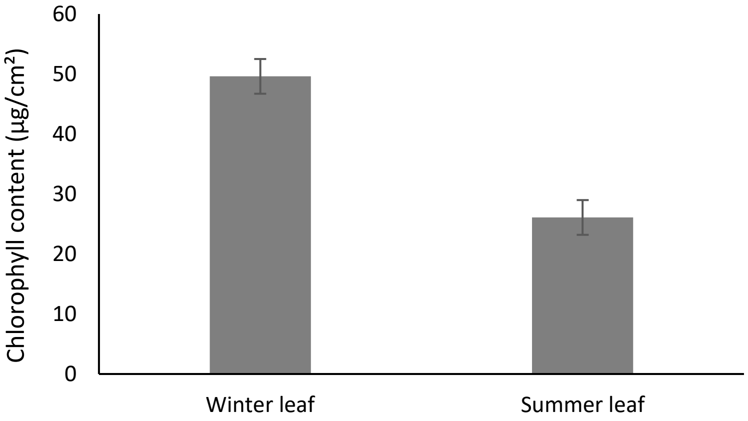
